# Quantifying a Frequency Modulation Response Biomarker in Responsive Neurostimulation

**DOI:** 10.1101/2020.12.17.423339

**Authors:** Praveen Venkatesh, Daniel Sneider, Mohammed Danish, Nathaniel D. Sisterson, Naoir Zaher, Alexandra Urban, Pulkit Grover, R. Mark Richardson, Vasileios Kokkinos

## Abstract

**Objective:** Responsive Neurostimulation (RNS) is an effective treatment for controlling seizures in patients with drug-resistant focal epilepsy who are not suitable candidates for resection surgery. A lack of tools for detecting and characterizing potential response biomarkers, however, contributes to a limited understanding of mechanisms by which RNS improves seizure control. We developed a method to quantify ictal frequency modulation, previously identified as a biomarker of clinical responsiveness to RNS.

**Approach:** Frequency modulation is characterized by shifts in power across spectral bands during ictal events, over several months of neurostimulation. This effect was quantified by partitioning each seizure pattern into segments with distinct spectral content and measuring the extent change from the baseline distribution of spectral content using the squared Earthmover’s distance.

**Main results:** We analyzed intracranial electroencephalography data from 13 patients who received RNS therapy, six of whom exhibited frequency modulation on expert evaluation. Subjects in the frequency modulation group had, on average, significantly larger and more sustained changes in their Earthmover’s distances (mean = 13.97×10^-3^ ± 1.197×10^-3^). In contrast, those subjects without expert-identified frequency modulation exhibited statistically insignificant or negligible distances (mean = 4.994×10^-3^ ± 0.732×10^-3^).

**Significance:** This method is the first step towards a quantitative, feedback-driven system for systematically optimizing RNS stimulation parameters, with an ultimate goal of truly personalized closed-loop therapy for epilepsy.

## 1. Introduction

The Responsive Neurostimulation (RNS) System is a closed-loop brain stimulation system, FDA-approved as an alternative treatment for drug-refractory focal epilepsy patients, who are not considered suitable candidates for resective surgery. The responsive neurostimulator automatically analyzes the intracranial electroencephalogram (iEEG), detects seizures and rapidly delivers electrical stimulation to suppress seizure activity (Kossof et al., 2004). Studies of Class 1 evidence have reported 44% seizure reduction at 1 year post-implantation, 53% at 2 years (Heck et al., 2014), and a 48-66% reduction in seizure occurrence between the 3rd and 6th post-implantation years in open-label continuation studies (Bergey et al., 2015). A median 70% of patients with both mesio-temporal and neocortical seizure onset experienced significant reduction in seizure frequency at 6 years post-implantation, 26-29% benefited from a post-implantation seizure-free period of at least 6 months and 15% experienced 1 year or longer free from seizures (Geller et al., 2017; Jobst et al., 2017). A more recent retrospective study reported a median reduction of 67% in seizures at 1 year, 75% at 2 years, 82% at ≥ 3 years and 74% at the most recent follow-up (Razavi et al., 2020). A prospective open label trial to evaluate RNS efficacy found a median 75% of patients experienced seizure reduction at 9 years post-implantation, and 35% of patients had ≥ 90% reduction in seizure frequency (Nair et al., 2020). These results compare favorably to alternate neuromodulation strategies such as vagal nerve stimulation and deep brain stimulation (Sisterson and Kokkinos, 2020).

Although the RNS System has been shown to provide improved seizure control and quality of life in patients with pharmaco-resistant focal epilepsy, its mechanisms of action are still under investigation. Historically, the primary hypothesis has been that patients experience a decreased seizure burden as a result of direct inhibition of ongoing ictal activity by electrical stimulation (Lesser et al., 1999; Kossoff et al., 2004; Skarpaas and Morrell, 2009; Morrell and Halpern, 2016). Although isolated samples of recordings and corresponding spectrograms supporting this hypothesis have been presented sporadically in the literature (Skarpaas and Morrell, 2009; Thomas and Jobst, 2015; Geller et al., 2017; Jobst et al., 2017), no systematic studies have presented an in-depth analysis of the brain’s response to closed-loop stimulation events. We recently tested the hypothesis of whether clinical efficacy arises from successful detection-triggered electrical stimulation and subsequent direct termination of seizure activity, and instead found evidence for an altogether different therapeutic mechanism involving chronic effects (Kokkinos et al., 2019). We evaluated time and time-frequency features beyond a narrow direct stimulation interval, throughout the time-course of the iEEG recordings, and discovered discrete categories of modulation effects associated with improved clinical outcomes that appeared independent of acute stimulation events. In contrast, detection-triggered stimulation effects were not associated with responsiveness to neurostimulation therapy. The identification of these biomarkers suggested that chronic responsive stimulation progressively disrupts the connectivity of the epileptogenic network and reduces the core synchronized population strength.

This prior work was largely qualitative, demonstrating visually appreciable changes in the time and frequency domains, similar to clinical observational practice, without a strong quantitative backbone. In the current study, we take one step further by developing quantitative metrics for describing iEEG response biomarkers in RNS recordings, focusing on the ictal frequency modulation effect.

## 2. Methods

### 2.1. Patients and Data

Data were obtained from an IRB-approved database of all epilepsy surgeries performed in a single surgeon practice at an academic medical center. All patients were diagnosed with refractory focal epilepsy according to current ILAE criteria (Berg et al., 2010; Fisher et al., 2017) and chronically implanted with closed-loop neurostimulation (RNS, NeuroPace, Mountain View, CA, USA) after consensus recommendation in a multidisciplinary epilepsy patient management conference. Data from 13 consecutive patients implanted with the RNS System between January 2015 and June 2018 were included in this study.

All patients had either neocortical or hippocampal implantation (Table 1), with two leads. Immediately post-implantation, the device was set to passive recording mode for approximately one month during which no stimulation was delivered, constituting the baseline epoch. Once the baseline activity was reviewed, stimulation parameters were configured and activated by the clinical team. In turn, detection-triggered stimulation therapy parameters were periodically modified in subsequent clinic visits, based on a clinical evaluation of seizure control status. The time intervals between RNS parameter modifications, during which both detection and stimulation settings remained unchanged, are referred to as programming epochs.

**Table 1.** Patient data

| **Patient** | **Age** | **Gender** | **Implantation sites** | **Postop Months** | **# Seizure patterns** | **Frequency modulation effects** | **Outcome (Engel Scale)** |
| --- | --- | --- | --- | --- | --- | --- | --- |
| 1 | 29 | M | Cortical malformation | 12 | 29 | Indirect | IIB |
| 2 | 22 | M | Neocortex | 13 | 22 | Indirect | IIB |
| 3 | 39 | F | Cortical malformation | 24 | 2,032 | Indirect | IIIA |
| 4 | 24 | M | Neocortex | 6 | 92 | Indirect | IIIA |
| 5 | 34 | F | Neocortex | 34 | 430 | Indirect | IB |
| 6 | 42 | F | Neocortex | 37 | 823 | Direct | IVB |
| 7 | 22 | F | Hippocampus | 31 | 487 | None | IVB |
| 8 | 47 | F | Cortical malformation | 41 | 263 | None | IIIA |
| 9 | 37 | M | Neocortex | 40 | 20 | None | IIA |
| 10 | 22 | F | Hippocampus | 30 | 1,013 | None | IVA |
| 11 | 30 | M | Hippocampus | 4 | 48 | None | IA |
| 12 | 39 | F | Hippocampus | 18 | 141 | None | IVB |
| 13 | 63 | F | Neocortex | 39 | 2,072 | None | IB |

iEEG recordings and additional metadata, including recording and detection settings, were retrieved from the NeuroPace Patient Data Management System (PDMS) using automated in-house custom-built software (Sisterson et al., 2019). All recordings were 90s periods of 4-channel iEEG, online band-pass filtered at 4-125Hz, sampled at 250Hz, and digitized by a 10-bit ADC. iEEG channels were recorded in a bipolar montage between neighboring electrode contacts, with the case of the RNS pulse generator acting as the amplifier ground. All electrode impedances measured below 1 kOhm in all recordings.

iEEG seizure pattern onsets were annotated based on visual evaluations conducted independently by an experienced epilepsy surgery neurophysiologist (VK) and a board-certified epileptologist (NZ). Inter-rater agreement was 99.8%. iEEG seizure pattern onset was defined as the point in time after which the iEEG background was no longer interictal and was followed by paroxysmal discharges with ictal features and morphology developing over time. The interictal background, both in the awake and sleep states, was appreciated from scheduled recordings and iEEGs that did not contain seizure patterns.

### 2.2. Preprocessing

The data corresponding to each programming epoch were comprised of a number of distinct files containing 90s of continuous iEEG data. These files were not always contiguous and therefore were processed independently of each other. Only seizure patterns that had an onset within a file were considered; seizure patterns for which the onset was not captured were discarded. For each seizure pattern, only the recording period between the onset and the end of the respective iEEG file was considered for analysis.

Beyond the baseline epoch, therapeutic neurostimulation was active for all epochs. Stimulation intervals appeared in the iEEG recording as an artifact, characterized by a flat response, followed by a brief burst with an exponential decay (we attribute the burst to charge being drawn back into the electrode as it switched from stimulation to recording). To remove this artifact, we searched for segments of iEEG data that were constant for at least 250ms across all channels and then eliminated these segments. We also fit an exponential curve to identify the subsequent burst, and disregarded data until it fell to 5% of the peak of the burst.

### 2.3. Processing: Spectral Content-based Partitioning of Seizure Patterns

We first used an algorithm to temporally partition each seizure pattern into segments based on their spectral content. For each patient, a single RNS channel was designated the onset electrode by visual inspection (Figure 1a). This channel was used for all subsequent analyses for that patient. The partitioning of iEEG seizure patterns comprised five discrete steps:

1. For each seizure pattern, we computed the Short-Time Fourier Transform (STFT), also known as the spectrogram, using a 1s-wide Kaiser window (*β* = 10) and a step size of one-sixteenth of a second (Figure 1b).
2. The spectrogram was thresholded using a 1s-wide moving window at steps of 0.25s. In each window, we thresholded all frequency values that had an amplitude less than half the peak amplitude within that window (Figure 1c).
3. We then analyzed the spectrogram for change points at intervals of 0.05s. For each putative change point, we compared the 2s window before that time instant to the 2s window following it. These two windows were compared for six distinct frequency bins (each 10Hz wide, spanning 0-60Hz). For each frequency bin, we performed a two-sample Kolmogorov-Smirnov (KS) test (Kass et al., 2014) to assess whether the distribution of frequency magnitudes in the two windows differed significantly. Accordingly, we computed six different p-values of the KS test for each frequency bin (Figure 1d).
4. The p-values were combined across bins using Fisher’s method to produce a single aggregate p-value at each tested time instant. It should be noted that this is not a statistically rigorous use of Fisher’s method, since the p-values being combined are, in general, not independent. However, this was found to be a reasonable heuristic for the purposes of segmentation (Figure 1e).
5. Finally, change points were declared using a peak-detection algorithm (Virtanen et al., 2020) on the negative logarithm of the aggregate p-values. We ensured that peaks had a prominence of at least “2 units”, implying that only p-values less than 0.01 were selected (Figure 1f, g).

**Figure 1.**
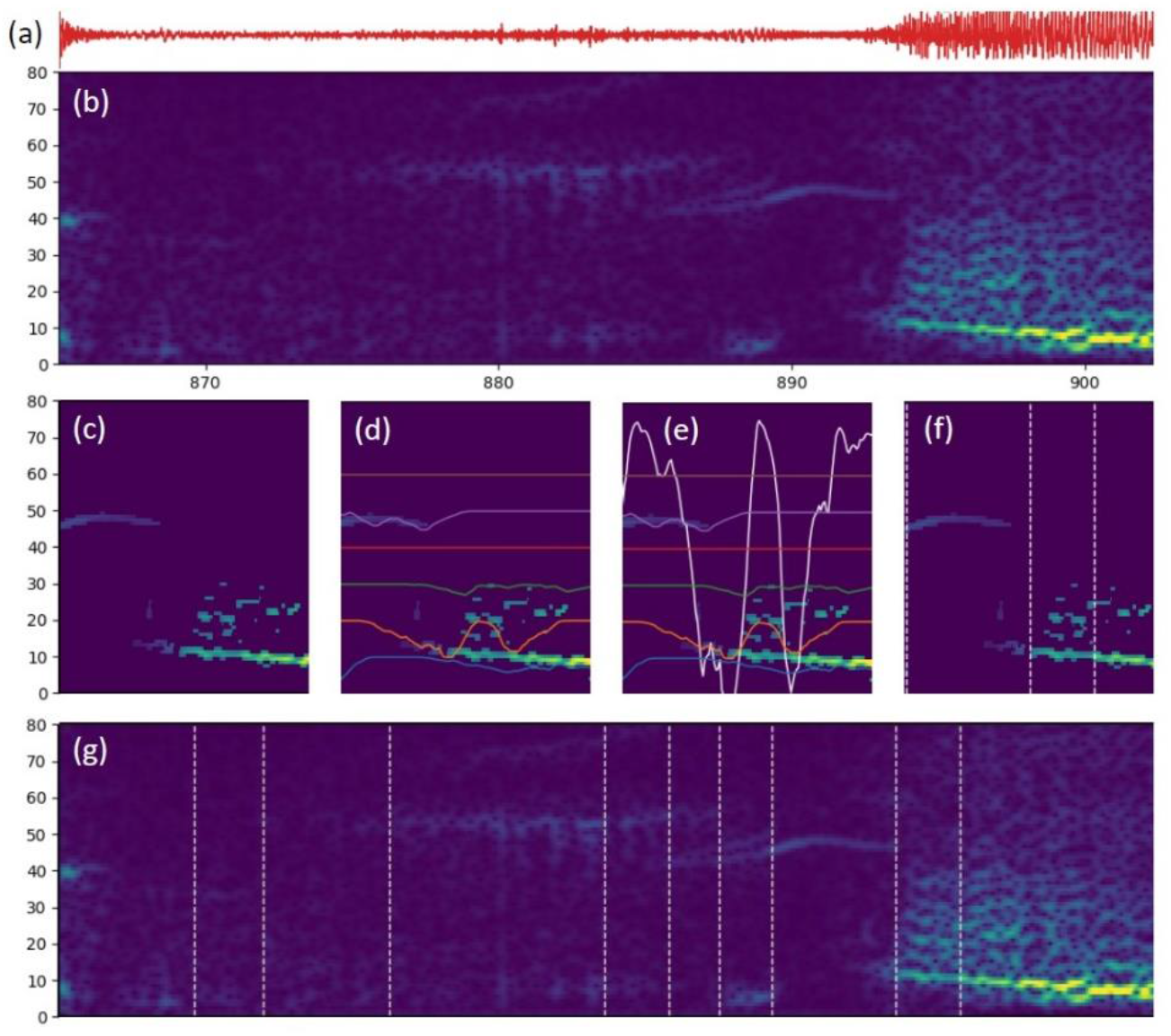
A depiction of the process of seizure partitioning. (a) Data from a single iEEG channel, here, the first 35s of a typical seizure from Patient 7 is shown for illustration. (b) The STFT of the aforementioned time-domain data. (c) STFT snippet after thresholding has been applied. (d) KS-test p-values for change detection superimposed upon the STFT snippet: each colored line represents log-scaled p-values for KS tests conducted for a 10Hz-wide frequency bin. For example, the orange line represents the 10-20Hz bin with the y-axis is scaled as follows: 20Hz corresponds to a p-value of 1, while 10Hz corresponds to a p-value of 10^-8^. The other lines are shifted according to their respective frequency bin and are scaled similarly. (e) The Fisher-combined p-value (after log-scaling) superimposed in white, with the y-axis scaled as follows: 80Hz corresponds to a combined p-value of 1, while 0Hz corresponds to a combined p-value of 10^-8^. (f) Peak detection applied to the combined p-values to detect change points, shown by white dashed vertical lines. (g) All change points for the first 35s of this seizure: note how points at which the seizure frequency content changes are correctly identified as change-points.

Of these steps, the segmentation algorithm was found to be particularly sensitive to the length of the thresholding window and the extent of thresholding. Other parameters did not affect the segmentation output considerably.

### 2.4. Processing: Quantification of Frequency Modulation in Seizure Patterns

Having partitioned each seizure pattern into segments based on frequency content, we proceeded to use these segments as features for quantifying frequency modulation in each patient. This process consisted of seven discrete steps:

1. We first condensed each segment into a three-dimensional vector, comprising the average frequency magnitude in three different frequency bands: 0-10Hz, 10-30Hz and 30-60Hz (Figure 2a, b).
2. Since each patient had different RNS signal amplitudes, we normalized each vector by its sum. This had the effect of balancing different patients’ signal energies and provided a common platform to compare the extent of frequency modulation across patients. Note that this step further condensed each segment to a two-dimensional vector lying on the standard simplex (Figure 2c).
3. We then grouped all segments arising from seizure patterns within a given programming epoch (indicated by colors in Figures 2b, c). Our quantification of frequency modulation depended only on the statistics of these segments across epochs. This ensured that we could reliably capture indirect frequency modulation, which was a result of chronic stimulation, and made us less likely to capture direct frequency modulation, which was an instantaneous result of stimulation (for details, refer to Kokkinos et al., 2019).
4. For each programming epoch, we computed the weighted empirical distribution of segments on the standard two-dimensional simplex (Figure 2d). Weights were assigned based on the time-span of each seizure segment. These distributions, therefore, captured the extent of variation in frequency amplitudes (weighted by duration) across all seizure patterns within each programming epoch.
5. The extent of frequency modulation between two epochs was quantified using a squared Earthmover’s distance (also called the Wasserstein distance; Villani, 2008) between their corresponding empirical distributions (Figure 2e). Intuitively, if we visualize these distributions as mounds of earth, then the Earthmover’s distance quantifies the minimum amount of “work” it would take to move the earth from one distribution so as to make it equal to the other. We computed this distance using an Optimal Transport algorithm (Flamary and Courty, 2017).
6. Finally, for every pair of epochs, we tested whether the computed distance was significantly different from zero by using a permutation test (Kass et al. 2014). We constructed an empirical null distribution by randomly permuting segments between the two epochs 10,000 times (keeping the number of segments in each epoch fixed) and computing the squared Earthmover’s distance between the resulting weighted empirical distributions each time. The estimated p-value was then the probability, under the empirical null distribution, of exceeding the true squared Earthmover’s distance.
7. Since we have multiple hypothesis tests for each patient, we identified all pairwise Earthmover’s distances that were significant at a *family-wise error rate* of 0.01 for each patient using Bonferroni’s method (Kass et al. 2014). This is equivalent to performing a Bonferroni correction against the number of pairwise comparisons for each patient (Figure 2f).

**Figure 2.**
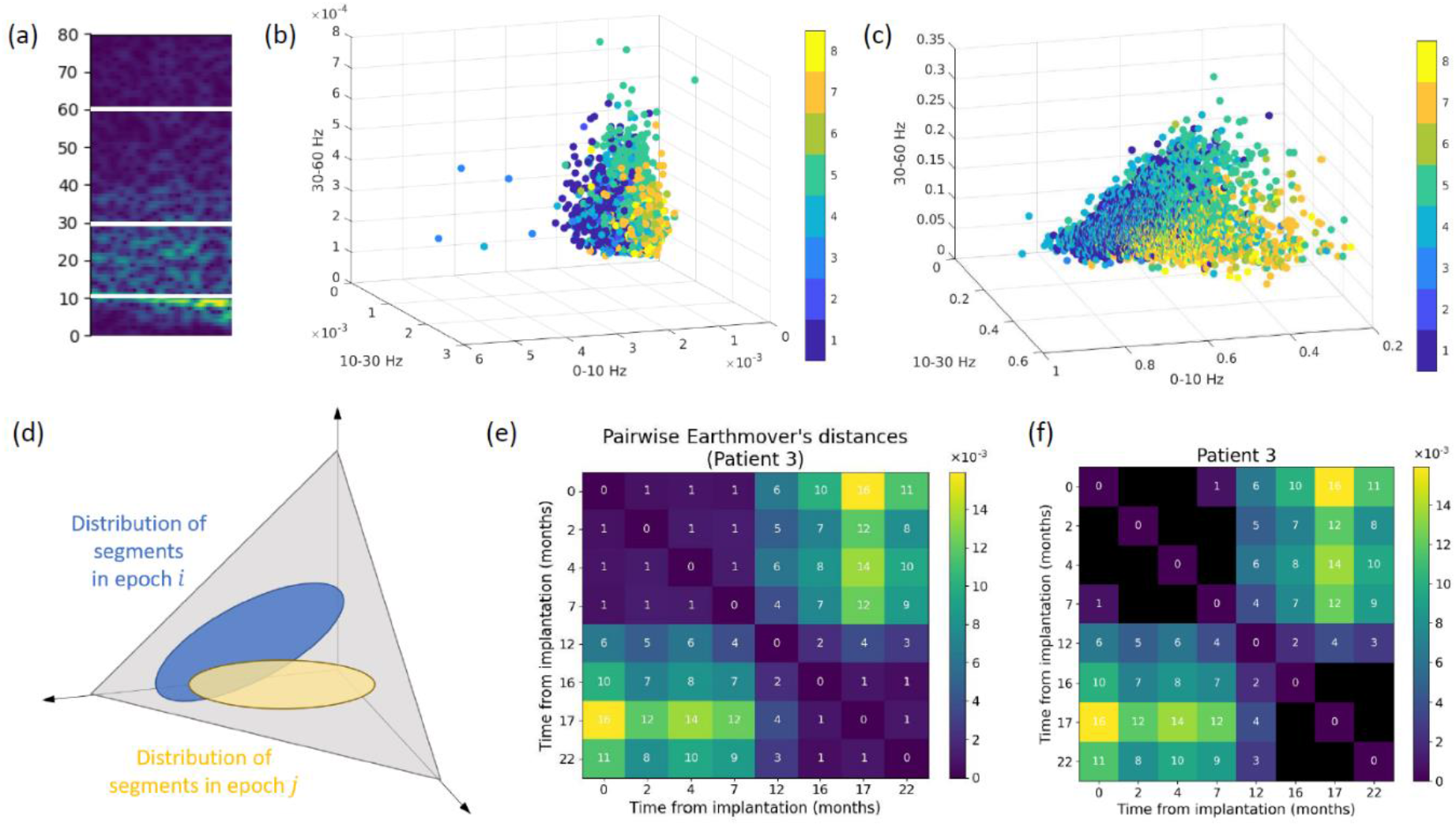
A depiction of the process of quantifying frequency modulation in a patient. (a) Each seizure segment is divided into three frequency bins: 0-10Hz, 10-30Hz and 30-60Hz. The average frequency magnitude in each of these bins becomes the axes of the 3D image in (b); each seizure segment is thus condensed to a single point. Figure (b) shows segments from all seizures across all epochs; the color of each segment represents the index of the programming epoch it appears in (data from Patient 3 shown for illustration). (c) Seizure segments normalized to have unit L1-norm. (d) A caricature of the distributions of segments from two different epochs; the squared Earthmover’s distance is computed between every such pair of empirical distributions, as shown in (e). Significance is evaluated using a permutation test and we report all distances significant at a family-wise error rate of 1% (for each patient), shown in (f).

### 2.5. Patient outcomes

Seizure outcomes were derived by personal impact of epilepsy scale (PIES) questionnaires (Fisher et al., 2015). PIES questionnaires were supplemented with three queries regarding seizure manifestation: (i) the estimated mean frequency of seizure occurrence before and after RNS implantation, per month in absolute numbers; (ii) the estimated mean severity of seizures, on a scale of 1 to 5 (1: not severe, 5: very severe); and (iii) the mean duration of seizures, in minutes. (Table 1). We used the Engel classification (Engel et al., 1993) to group our patients as either responders (Engel class III or better) or non-responders (Engel class IV), based on the scores of the 3 seizure manifestation variables.

## 3. Results

We analyzed data from 13 patients implanted with the RNS System (5 male, mean age = 34 years, age range 22-63). The mean time after implantation to activation of responsive stimulation was 6 weeks (SD: 3.6). A total of 23,868 iEEG files were visually reviewed, corresponding to 316 months of post-surgical implantation recordings spanning a 41-month total study period. 7,472 seizure patterns were identified. After analysis presented elsewhere (Kokkinos et al., 2019) and extended in this cohort of patients, both direct and indirect ictal frequency modulation was appreciated in six of our patients (patients 1 through 6), while the remaining seven (patients 7 through 13) did not demonstrate any visually appreciable change in ictal frequency content.

For each patient’s recordings, we measured the extent of frequency modulation between every pair of programming epochs by using a squared Earthmover’s distance between the weighted empirical distributions of the epochs’ seizure segments. Recordings from patients in the frequency modulation group, on average, exhibited larger magnitudes of squared Earthmover’s distances (mean = 13.97×10^-3^ ± 1.197×10^-3^ among statistically significant values reported in Figures 3 and 4 combined, for Patients 1-6; mean = 14.49×10^-3^ ± 1.282×10^-3^ among statistically significant values reported in Figure 3 alone, for Patients 3-6) compared to recordings from patients that did not exhibit frequency modulation (mean = 4.994×10^-3^ ± 0.732×10^-3^ among statistically significant values reported in Figure 5 for Patients 7-13).

**Figure 3.**
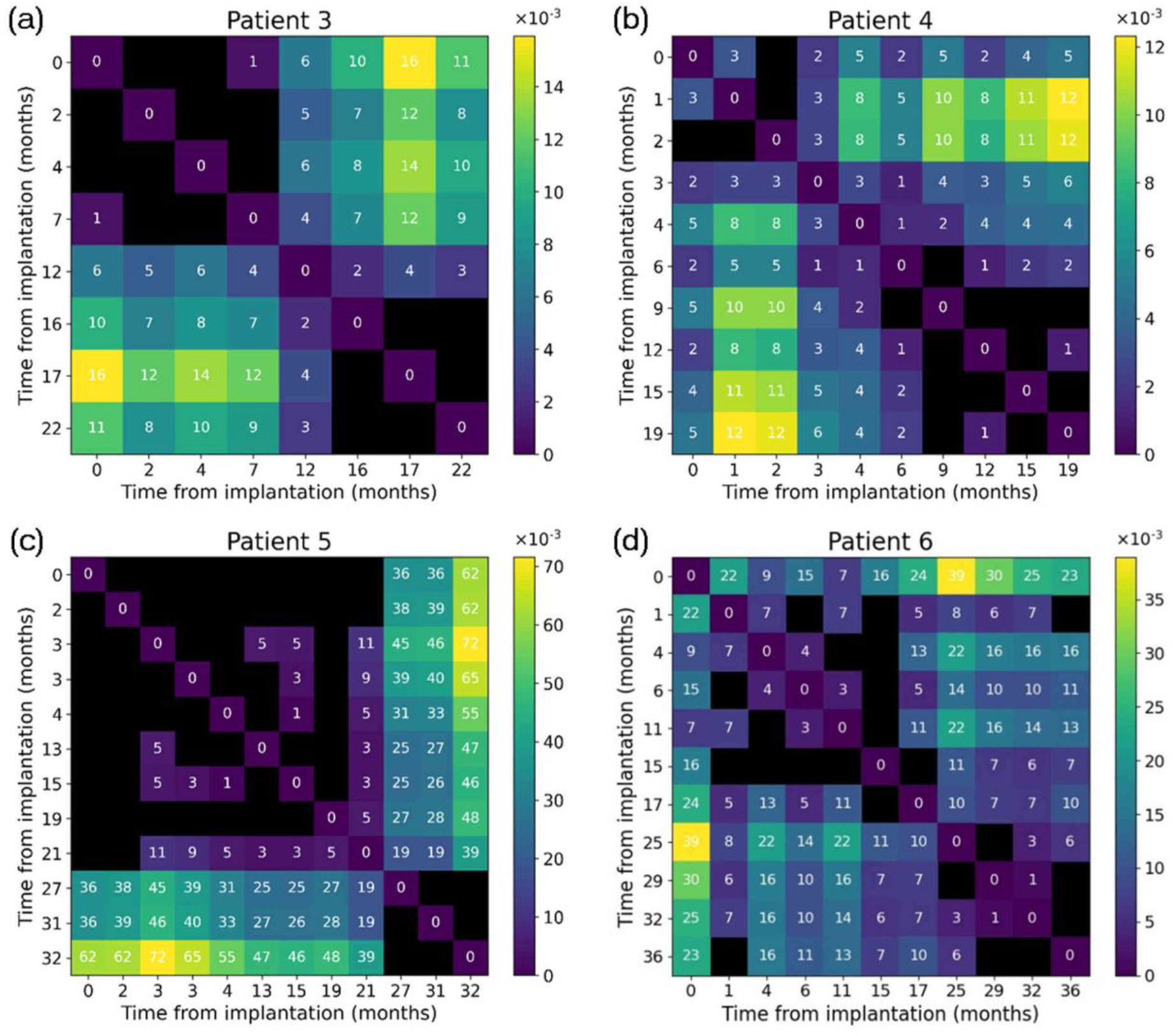
Earthmover’s distances between every pair of programming epochs, in patients who showed indirect frequency modulation according to an expert visual evaluation.

**Figure 4.**
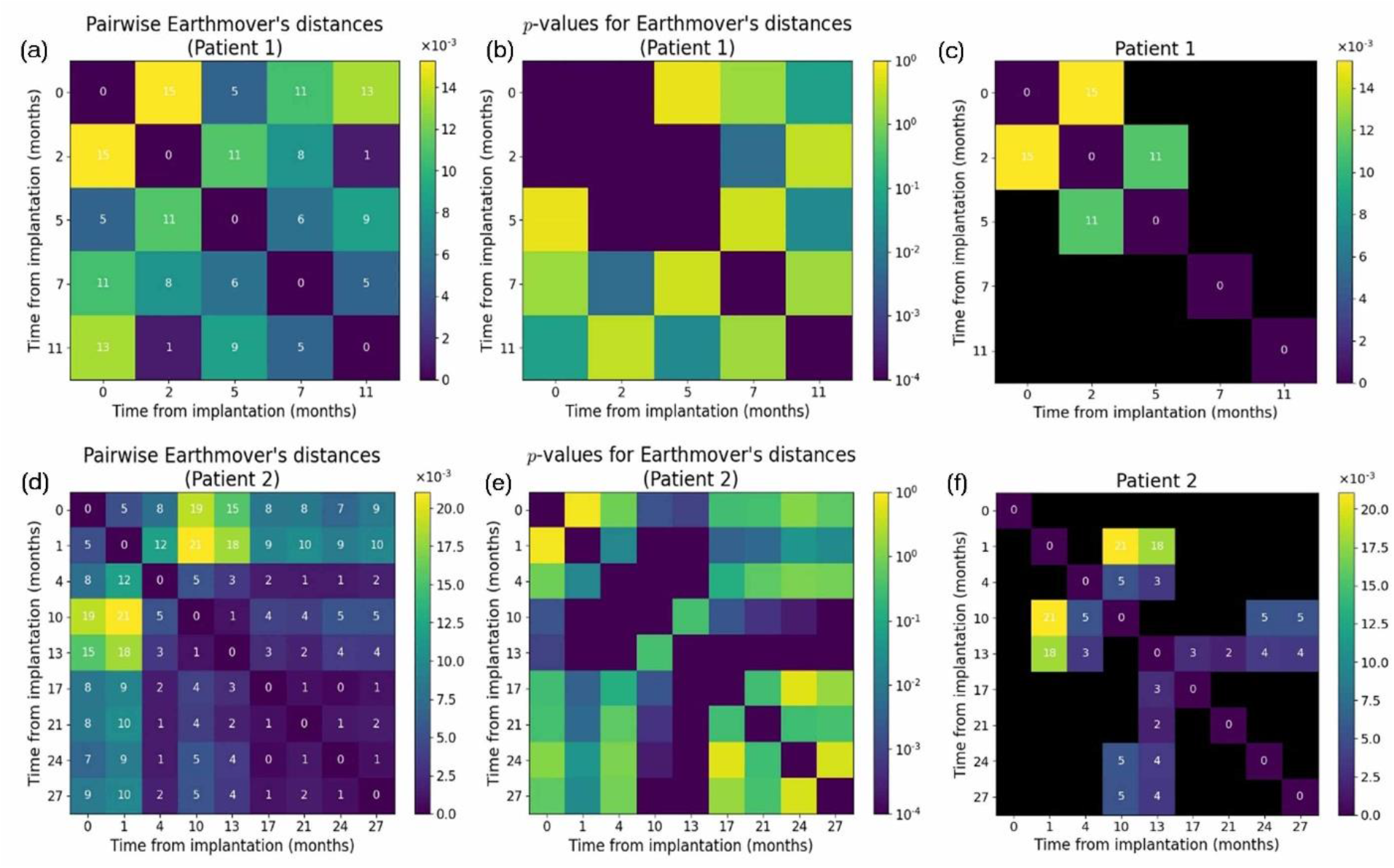
Results for patients 1 and 2, both of whom showed frequency modulation in an expert evaluation. (a-c) Patient 1 had too few seizures to be able to see frequency modulation when using our measure. (d-f) Patient 2 had too few seizures in the first few epochs, which is where the squared Earthmover’s distances are more pronounced. Thus, in both cases, statistical significance of these distances could not be established.

Analysis of recordings in the frequency modulation group produced Earthmover’s distance matrices with a block-diagonal structure, with larger distances in off-diagonal blocks and distances close to zero on diagonal blocks (Figure 3). These recordings followed a pattern: Earthmover’s distances show a marked increase after a specific programming epoch that was subsequently sustained. This finding is represented by a distinct block-matrix structure: diagonal blocks appear dark, having either very small or statistically insignificant distances, while off-diagonal blocks are bright, indicating a sustained change in the frequency content of these patients’ seizure patterns. For example, in Patient 3, the squared Earthmover’s distance from the baseline epoch remains close to zero (or indistinguishable from zero) for the first 12 months after implantation; subsequently it rises to an average value of 10.75×10^-3^ over the next 10 months (Figure 3a). Similarly, recordings from Patient 5 showed squared Earthmover’s distances indistinguishable from zero until the 27^th^ month post-implantation, after which it rises in the last three epochs, attaining a maximum of 72×10^-3^ (the largest distance observed within our cohort) (Figure 3c). Recordings from Patient 6 were classified as showing direct frequency modulation in the expert evaluation. However, the analysis demonstrated large, statistically significant changes in Earthmover’s distance, as well as the visually apparent block-diagonal structure, suggesting that the method is sensitive to both indirect and indirect frequency modulation effects.

Patients 1 and 2 were assigned to the frequency modulation group, however, the programming epochs that contributed most to frequency modulation had an insufficient number of seizure patterns to obtain statistically meaningful Earthmover’s distances (Figure 4). Patient 1 had too few seizure events: 2, 18, 5, 4 and 9 events, respectively, in their five programming epochs (Figure 4a-c). Patient 2 showed signs of frequency modulation, as is apparent from the box-diagonal structure in the full matrix (Figure 4d-f), but the epochs that appeared to show the largest modulation effects were also those with the smallest number of seizure patterns (the first five programming epochs had 2, 5, 12, 0 and 15 recorded seizure events; an epoch without seizure events was not included in the matrix). Thus, several distances did not pass the statistical threshold at a family-wise error rate of 0.01. Based on the data available to us, we find that our algorithm starts becoming effective when working with at least 15-20 seizure events per programming epoch and performs more reliably as the number of seizure patterns per programming epoch increases.

Recordings from the group without ictal frequency modulation did not demonstrate a clear pattern of squared Earthmover’s distances change and often had distances that were either small or statistically insignificant (Figure 5). For example, in recordings from Patient 8, almost all squared Earthmover’s distances are statistically insignificant, the remaining few being small relative to those values from the group of patients who exhibited statistically significant frequency modulation. In the case of Patient 7, most distances were significant, however they were all quite small (< 6×10^-3^) and showed no clear block-diagonal structure.

**Figure 5.**
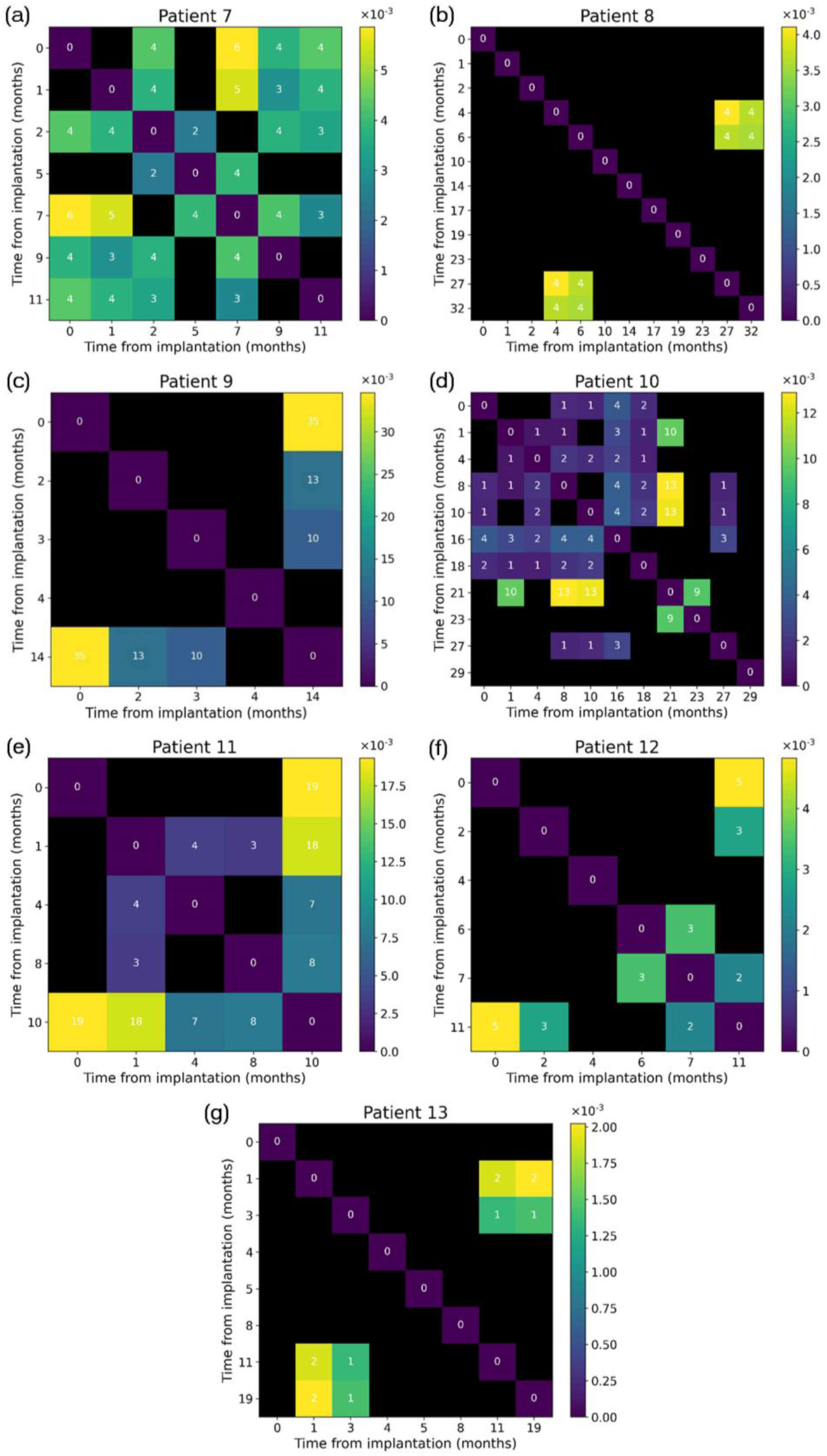
Earthmover’s distances between every pair of programming epochs in patients who did not exhibit indirect frequency modulation effects according to an expert visual evaluation.

## 4. Discussion

Understanding the effects of responsive neurostimulation in any given epilepsy patient requires substantial effort by clinicians to manually review recorded electrocorticograms. As new potential biomarkers of clinical response are discovered, the burden on clinicians to recognize and quantify these electrophysiological responses will continuously increase. It is imperative, therefore, to develop tools to automate the process of response biomarker detection. In this study we described a method that differentiated patients having recordings in which the ictal frequency modulation response biomarker was present. These recordings exhibited Earthmover’s distance matrices with a block-diagonal structure, with larger distances in off-diagonal blocks and small or statistically non-significant distances on diagonal blocks. In contrast, recordings from patients in which no frequency modulation effect was present exhibited no clear matrix pattern, and often had squared Earthmover’s distances that are either uniformly low or statistically non-significant. We also found that the Earthmover’s assay for quantifying frequency modulation performed most reliably when the number of seizure events increased to at least 15-20 per programming epoch.

The efficacy of electrical stimulation in seizure control has been clearly demonstrated throughout the literature. Penfield was the first to observe and report the inhibitory effects induced by electrical stimulation on active epileptogenic neural tissue (Penfield and Jasper, 1954). Since then, cortical electrical stimulation was found repeatedly to have a significant inhibitory effect on interictal and ictal cortical activity in epilepsy patients (Velasco et al., 2000; Yamamoto et al., 2002; Kinoshita et al., 2004; Kossoff et al., 2005; Kinoshita et al., 2005; Yamamoto et al, 2006; Elisevich et al., 2006; Velasco et al., 2009; Child et al., 2014; Lundstrom et al., 2016, 2017; Valentín et al., 2017; Kerezoudis et al., 2017). The RNS system became a valuable surgical option for patients refractory to both anti-epileptic drugs and traditional epilepsy surgery (Stacey and Litt, 2008), with data supporting its superiority to medical management (Heck et al., 2014; Geller et al., 2017; Jobst et al., 2017; Razavi et al., 2020; Nair et al., 2020). However, apart from a few original publications (Lesser et al., 1999; Kossoff et al., 2004; Osorio et al., 2005), little was known about how closed-loop stimulation affects the time-course of electrographic seizure patterns or its mechanisms of action for reducing seizure severity and frequency until recently. Previously, the mechanism was assumed to be direct, wherein the application of an electric pulse close to the origin of the electrographic seizure pattern interrupts its evolution and returns the iEEG background to its interictal state (Kossoff et al., 2004; Skarpaas and Morrell, 2009; Morrell and Halpern, 2016).

In our previous work, we highlighted the existence of indirect modulation effects that correlated with responsiveness to RNS, thereby providing a novel paradigm for RNS’s mechanism of action (Kokkinos et al., 2019). In particular, the presence of frequency modulation effects among our earlier findings showed that parts of the underlying epileptogenic tissue can be progressively retuned by stimulation to oscillate at different frequencies than before responsive neurostimulation. These findings were consistent with the reported increase of patient responsiveness as a function of time (Bergey et al., 2015; Razavi et al., 2020). Our initial study was largely qualitative, adhering to the principles of routine clinical evaluation of iEEG recordings. In the present study, we took one step further towards quantifying and reducing the subjectivity of our observations on ictal frequency modulation in RNS.

There are currently no patient-specific guidelines for optimal regulation of either detection or stimulation parameters for RNS (Sisterson et al., 2019; 2020). Quantification of the frequency modulation biomarker describe here, however, provides insight for understanding how this type of information might be used prospectively to optimize stimulation parameters. Take the example of data from non-responder Patient 8 shown in Figure 5b, which did not demonstrate visually appreciable frequency modulation. The majority of programming epochs demonstrated no sign of frequency modulation. However, in programming epochs corresponding to months 4-6 post-implantation, statistically significant indications of frequency modulation appear. It is possible that the specific combination of detection and stimulation settings during that treatment phase had the potential to modulate the epileptogenic neuronal substrate and change the ictal oscillation frequency range, without rendering these changes visually appreciable. If the clinician had received this information at the time, settings might have been preserved and the patient’s matrix might have developed like that of responder Patient 5 in Figure 3c, who demonstrated frequency modulation effects only after 27 months post-implantation and a considerable number of changes in settings. Our results show that our method not only quantifies specific modulation effects but can also act as a pointer towards the optimal settings that provoke said neuromodulation effects.

## 5. Conclusion

We developed a method for quantifying indirect ictal frequency modulation in patients undergoing RNS treatment. To our knowledge, this is the first metric that quantifies a biomarker of the longterm efficacy of responsive neurostimulation. Given enough samples of seizure patterns, this method demonstrated the potential to match qualitative expert assessments and potentially to predict long-term clinical outcomes. Statistically significant fluctuations in the Earthmover’s metric of patients who did not show consistent indirect frequency modulation may indicate the direction in which RNS stimulation parameters must be tuned to produce this effect and improve seizure control. Quantification of indirect frequency modulation is but a first step in identifying and quantifying biomarkers of the long-term effects of RNS, as several other biomarkers have been previously identified and discussed qualitatively (Kokkinos et al., 2019; Sisterson and Kokkinos, 2020), with the long-term goal of achieving genuinely personalized closed-loop brain modulation for epilepsy.

## Acknowledgments

P.V. was supported in part by a Dowd Fellowship from the College of Engineering at Carnegie Mellon University, and in part by a Fellowship in Digital Health from the Center for Machine Learning and Health at Carnegie Mellon University. The authors would like to thank Philip and Marsha Dowd for their financial support and encouragement.

